# Diurnal fluctuations in steroid hormones tied to variation in intrinsic functional connectivity in a densely sampled male

**DOI:** 10.1101/2023.10.16.562607

**Authors:** Hannah Grotzinger, Laura Pritschet, Pavel Shapturenka, Tyler Santander, Elle Murata, Emily G. Jacobs

## Abstract

Most of mammalian physiology is under the control of biological rhythms, including the endocrine system with time-varying hormone secretion. Precision neuroimaging studies provide unique insights into the means through which our endocrine system regulates dynamic properties of the human brain. Recently, we established estrogen’s ability to drive widespread patterns of connectivity and enhance the functional efficiency of large-scale brain networks in a woman sampled every 24h across 30 consecutive days, capturing a complete menstrual cycle. Steroid hormone production also follows a pronounced sinusoidal pattern, with a peak in testosterone between 6-7am and nadir between 7-8pm. To capture the brain’s response to diurnal changes in hormone production, we carried out a companion precision imaging study of a healthy adult man who completed MRI and venipuncture every 12-24 hours across 30 consecutive days. Results confirmed robust diurnal fluctuations in testosterone, cortisol, and estradiol. Standardized regression analyses revealed predominantly positive associations between testosterone, cortisol, and estradiol concentrations and whole-brain patterns of coherence. In particular, functional connectivity in Dorsal Attention and Salience/Ventral Attention Networks were coupled with diurnally fluctuating hormones. Further, comparing dense-sampling datasets between a man and naturally-cycling woman revealed that fluctuations in sex hormones are tied to patterns of whole-brain coherence to a comparable degree in both sexes. Together, these findings enhance our understanding of steroid hormones as rapid neuromodulators and provide evidence that diurnal changes in steroid hormones are tied to patterns of whole-brain functional connectivity.

**Significance Statement:** Diurnal variation is an essential biorhythm, yet the relationship between diurnal fluctuations in steroid hormones and the functional architecture of the human brain is virtually unknown. This precision neuroimaging study suggests that endogenous fluctuations in testosterone, estradiol, and cortisol concentrations are tied to rhythmic changes in coherence across the brain. Precision imaging studies that track individuals across major endocrine transitions (e.g. the diurnal cycle and menstrual cycle) demonstrate steroid hormones’ ability to modulate the functional architecture of the brain in both sexes, and provide a starting point for future studies to probe the functional significance of these rhythms for behavior.

---

The mammalian brain is densely packed with steroid hormone receptors, yet the extent to which these signaling molecules—including estrogen, testosterone, and cortisol—influence the large-scale functional architecture of the human brain is remarkably understudied. At the cellular level, steroid hormones regulate synaptic plasticity in the hippocampus and prefrontal cortex (PFC) (Galea et al., 2017; Taxier et al., 2020). At the behavioral level, hormonal transitions such as puberty (Brouwer et al., 2015; McDermott et al., 2012; Pattwell et al., 2013), the menstrual cycle (Pritschet et al., 2020; Taylor et al., 2020; Zsido et al., 2022), pregnancy (Carmona et al., 2019; Hoekzema et al., 2017), menopause (Jacobs et al., 2017), and andropause (Janowsky, 2006) are tied to changes in brain function and structure. Fluctuations in hormone concentrations across shorter timescales, like diurnal rhythms, may also influence brain function. Existing neuroimaging studies of circadian rhythms focus on external influences that impact the human circadian clock, including sleep (Fang & Rao, 2017; Frank et al., 2013; Jiang et al., 2016; Mong et al., 2011), exposure to light sources (Schoonderwoerd et al., 2022), and psychopathology (Chen et al., 2022; Frank et al., 2013; McKenna et al., 2014), but the fundamental relationship between diurnal variation in sex steroid hormones and the large scale functional organization of the human brain is virtually unknown.

Human brain imaging studies often draw inferences about hormone-brain relationships via cross-sectional designs that pool data across subjects sampled at a single timepoint. In other cases, “sparse-sampling” longitudinal designs track changes within individuals at discreet timepoints—for example, across phases of the menstrual cycle (Hjelmervik et al., 2014; Lisofsky et al., 2015; Protopopescu et al., 2008; Weis et al., 2008) or stages of the menopausal transition (Maki & Resnick, 2000; Mosconi et al., 2018). However, a central feature of the mammalian endocrine system is that hormone secretion varies over time. Cross-sectional studies that capture a snapshot of the brain at one timepoint (or one endocrine state) could obscure the full range of brain-hormone dynamics as they unfold across these sinusoidal rhythms. Neuroimaging studies that densely sample individuals over timescales of days, weeks, or even months (Fedorenko, 2021; Gordon et al., 2017; Poldrack et al., 2015) are now being leveraged to provide unique insights into the role our endocrine system plays in regulating the dynamic nature of the human brain over time (Jacobs, 2023; Pritschet et al., 2021). Precision imaging of the human brain across neuroendocrine transitions adds a novel methodological approach for understanding the influence of biological rhythms on the brain (Jacobs, 2023; Pritschet et al., 2020, 2021; Taylor et al., 2020; Fitzgerald et al., 2020; Mueller et al., 2021; Zsido et al., 2022; de Filippe et al., 2021).

Previously, we examined the relationship between sex hormones and functional brain networks in a woman sampled every 24h for 30 consecutive days (Pritschet et al., 2020, 2021; Fitzgerald et al., 2020; Mueller et al., 2021; Taylor et al., 2020). Across the menstrual cycle, rhythmic changes in 17-β estradiol drive increases in functional coherence and enhance global efficiency in several intrinsic brain networks, including the Default Mode, Dorsal Attention, and Temporal Parietal Networks. Notably, many of these network hubs are populated with sex hormone receptors. These findings provided a foundation for understanding estrogen-driven changes in large-scale brain network organization, but we lacked complementary data to test these associations in a densely-sampled man.

While naturally-cycling women often undergo rhythmic changes in hormone production over a ∼28 day reproductive cycle, men (and women) experience a diurnal cycle: testosterone concentrations peak in the morning and decline by ∼50% or more throughout the day (Barberia et al., 1973; Diver et al., 2003; Rose et al., 1972), with estradiol and cortisol production following suit (∼35% and ∼85%, respectively) (Ankarberg-Lindgren & Norjavaara, 2008; Berg & Wynne-Edwards, 2001). To determine the influence of these diurnal hormone fluctuations on functional brain networks, we used a dense-sampling design to collect MRI, serum, saliva, and mood data from a healthy adult man every 12-24h for 30 consecutive days. Based on our original findings in a naturally-cycling woman, we predicted that the Default Mode, Dorsal Attention, and Temporal Parietal Networks would remain sensitive to steroid hormones. Direct comparisons between datasets from the man and woman allowed us to examine whether the magnitude of brain-hormone associations is comparable by sex across the brain. Results demonstrate that diurnal changes in steroid hormones are associated with increased whole-brain functional connectivity in a densely sampled man. Day-to-day changes in testosterone, estradiol, and cortisol show widespread associations with cortical network dynamics, particularly Dorsal and Ventral Attention Networks. Finally, comparing dense-sampling datasets between the man and naturally-cycling woman reveals that fluctuations in estrogen are tied to patterns of whole-brain coherence to a comparable degree in both sexes. Together, findings from this study enhance our understanding of testosterone, cortisol, and estradiol as rapid neuromodulatory hormones, and provide evidence that diurnal changes in steroid hormone production impact the brain’s functional network architecture.

## Materials and Methods

### Participant

The participant was a 26-year-old right-handed Caucasian male with no history of neuropsychiatric diagnosis, endocrine disorders, or prior head trauma. The participant gave written informed consent for a study approved by the University of California, Santa Barbara Human Subjects Committee and was paid for their participation in the study.

### Experimental design

The methods for this study parallel those reported in Pritschet et al., 2020. The participant (author P.S.) underwent venipuncture and brain imaging every 12-24h for 30 consecutive days. At each session the participant completed a daily questionnaire (see *Behavioral Assessments*), followed by endocrine sampling at 7am (morning sessions) and at 8pm (evening sessions). The participant gave a 2mL saliva sample at each session, followed by a blood sample. On days with two sessions, the participant underwent one blood draw per day (**Fig. 1**) per safety guidelines.

**Figure 1.**
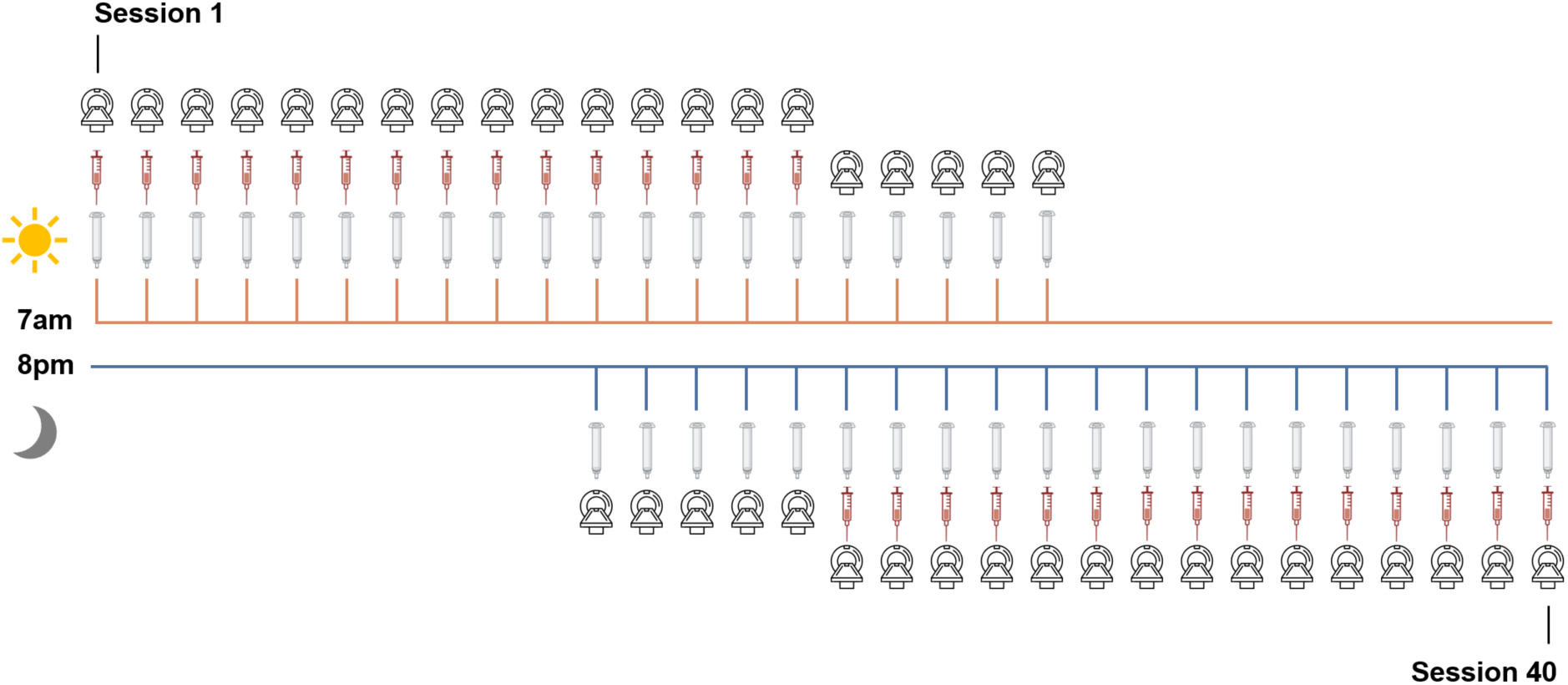
Sampling rate of MRI, venipuncture and saliva acquisition. Forty time-locked sessions were performed over 30 consecutive days; n=20 at 7am and n=20 at 8pm.

Morning endocrine samples were collected after at least 8 hours of overnight fasting, and evening endocrine samples were collected following an hour and a half of abstaining from consumption of food or drink (excluding water). The participant refrained from consuming caffeinated beverages before each morning session.

#### Behavioral assessments

The following scales (adapted to reflect the past 12-24h) were administered at each session during the daily survey: the Perceived Stress Scale (PSS) (Cohen et al., 1983), Pittsburgh Sleep Quality Index (PSQI) (Buysse et al., 1989), State-Trait Anxiety Inventory for Adults (STAI) (Speilberger & Vagg, 1984), Profile of Mood States (POMS) (Pollock et al., 1979), and Aggression Questionnaire (Buss & Perry, 1992). The questionnaire for the evening sessions excluded the PSQI to avoid redundancy. All mood measures fell within standard reference ranges. There were no associations between hormones and indices of sleep quality, stress, or anxiety (**Fig. 2**). After correcting for multiple comparisons (Bonferroni-corrected for 91 comparisons), cortisol concentrations were positively correlated with aggression (*r*(38) =0.54, *p*

**Figure 2.**
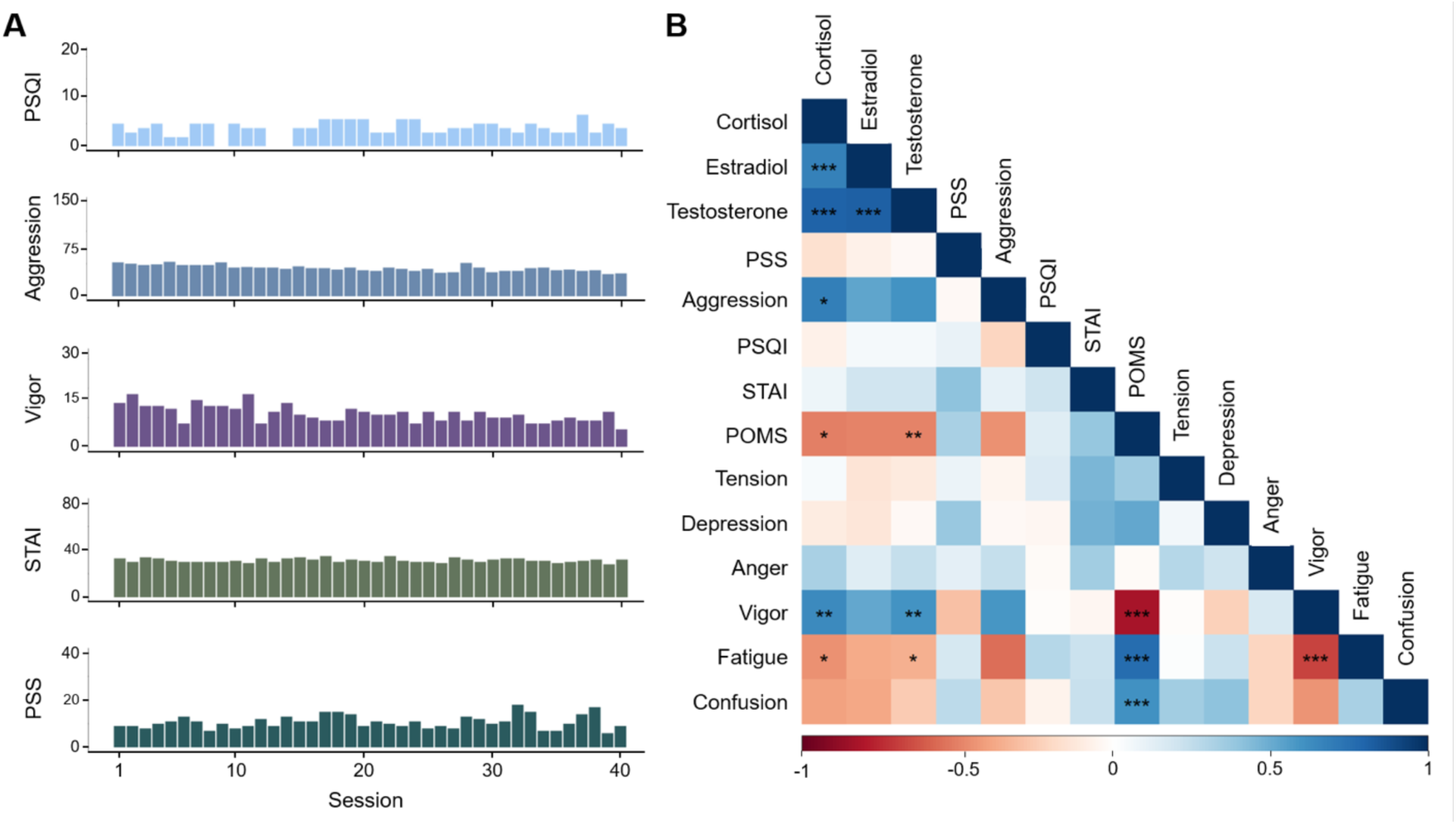
Survey measures were within the standard range and consistent across the study. **(A)** Measures of sleep, aggression, vigor, anxiety, and stress at each session. **(B)** Correlation plot depicts relationships between steroid hormones and mood. Cool colors indicate positive correlations, warm colors indicate negative correlations. Asterisks indicate significant correlations after Bonferroni correction (* *p* < 0.00055, ** *p* < 0.00011, *** *p* < 0.00001).

= .0003) and vigor (*r*(38) = 0.58, *p* = .0001), and negatively correlated with overall POMS scores (*r*(38) = -0.55, *p* = .0002) and fatigue (*r*(38) = -0.54, *p* = .0003). Total testosterone concentrations were also significantly positively correlated with vigor (*r*(38) = 0.58, *p* = .0001) and negatively correlated with overall POMS scores (*r*(38) = -0.58, *p* = .0001) and fatigue (*r*(38) = -0.53, *p* = .0005). Higher POMS scores indicate greater mood disturbance. Estradiol concentrations were not significantly correlated with any survey measures after Bonferroni correction.

#### Endocrine Procedures

A ∼2mL saliva sample was obtained via passive drooling into a wide mouthed plastic cryovial. The participant refrained from eating and drinking (besides water) at least 1.5 hours before saliva sample collection, and the morning samples were collected after fasting overnight. The participant pooled saliva for ∼5-10 minutes before depositing the sample into the cryovial to determine total testosterone and cortisol concentrations. The sample was stored at -20°C until assayed. Saliva concentrations were determined via enzyme immunoassay at Brigham and Women’s Hospital Research Assay Core.

Immediately after the saliva sample was obtained, a licensed phlebotomist inserted a saline-lock intravenous line into either the dominant or non-dominant hand or forearm daily to evaluate total testosterone, free testosterone, cortisol, and 17β-estradiol concentrations. One 10cc mL blood sample was collected in a vacutainer SST (BD Diagnostic Systems) each session. The sample clotted at room temperature for ∼60 min until centrifugation (2100 x g for 10 min) and was then aliquoted into three 2 ml microtubes. Serum samples were stored at -20°C until assayed. Serum concentrations were determined via liquid chromatography mass-spectrometry at the Brigham and Women’s Hospital Research Assay Core. Assay sensitivities, dynamic range, and intra-assay coefficients of variation (respectively) were as follows: estradiol, 1 pg/mL, 1– 500 pg/mL, < 5% relative standard deviation (*RSD*); testosterone, 1.0 ng/dL, 1–200 ng/dL, <2% *RSD*; cortisol, 0.5 ng/mL, 0.5-250 pg/mL, <8% *RSD*.

Note that estradiol and free testosterone measurements were acquired from serum samples, resulting in 30 timepoints.

### MRI acquisition

At each session, the participant underwent a structural MRI and 15-minute eyes-open resting-state scan conducted on a Siemens 3T Prisma scanner equipped with a 64-channel phased-array head coil. High-resolution anatomical scans were acquired using a T1-weighted magnetization prepared rapid gradient echo (MPRAGE) sequence (TR = 2500 ms, TE = 2.31 ms, TI = 934 ms, flip angle = 7°, .8 mm thickness), followed by a gradient echo fieldmap (TR = 758 ms, TE1 = 4.92 ms, TE2 = 7.38 ms, flip angle = 60°). Functional data were obtained via *T2\**-weighted multiband echo-planar imaging (EPI) sequence sensitive to blood oxygenation level-dependent (BOLD) contrast (72 oblique slices, TR = 720 ms, TE = 37 ms, voxel size = 2mm^3^, flip angle = 56°, MB factor = 8). In an effort to minimize motion, the head was secured with a 3D-printed foam head case. Minimal motion was detected throughout the experiment (**Fig. S1**).

### MRI preprocessing

#### Functional preprocessing

Initial preprocessing was performed using the Statistical Parametric Mapping 12 software (SPM12, Wellcome Trust Centre for Neuroimaging, London) in MATLAB. Functional data were realigned and unwarped to correct for head motion and geometric deformations due to motion and magnetic field inhomogeneities; the mean motion-corrected image was then coregistered to the high-resolution anatomical image. All scans were normalized to a subject-specific template using Advanced Normalization Tools’ (ANTs) multivariate template construction (Avants et al., 2011). A 4mm full-width at half-maximum (FWHM) isotropic Gaussian kernel was subsequently applied to smooth the functional data. Further processing was performed using in-house MATLAB scripts. Global signal scaling (median=1,000) was applied to account for fluctuations in signal intensity across space and time, and voxelwise timeseries were linearly detrended. Residual BOLD signal from each voxel was extracted after removing the effects of head motion and five physiological noise components (derived from CSF and white matter signal—although our use of coherence for functional connectivity allows for the estimation of frequency-specific covariances in spectral components below the range typically contaminated by physiological noise). Motion was modeled based on the Friston-24 approach, using a Volterra expansion of translational/rotational motion parameters, accounting for autoregressive and nonlinear effects of head motion on the BOLD signal (Friston et al., 1996). All nuisance regressors were detrended to match the BOLD timeseries. We note that steroid hormone concentrations were related to head motion (framewise displacement; FWD) due to generally greater movement during the evening sessions than morning sessions (*t*(32.15) = -3.85, *p* < .001). However, the average FWD was exceedingly minimal (*M* = 8 microns, *SD* = 4 microns), with a maximum of 930 microns across all 40 sessions. Nevertheless, to ensure this did not confound our findings, we specified a series of binary spike regressors for any frames that had framewise displacements greater than 500 microns (necessary in 17/40 sessions). To further ensure the robustness of our results, we analyzed the data with global signal regression (GSR) included, though analysis without global signal regression produced a near identical pattern of results (**Fig. S5**).

#### Functional connectivity estimation

Functional network nodes were defined by parcellating the brain based on the 400-region cortical (Schaefer et al., 2018) and 15 regions from the Harvard-Oxford subcortical atlas (http://www.fmrib.ox.ac.uk/fsl/). A summary timecourse for each session was extracted per node by taking the first eigenvariate across functional volumes (Friston et al., 2006). These regional timeseries were then decomposed into several frequency bands using a maximal overlap discrete wavelet transform. Low-frequency fluctuations in wavelets 3-6 (∼.01-.17 Hz) were selected for subsequent connectivity analyses (Patel & Bullmore, 2016). We estimated the *spectral* association between regional timeseries using magnitude-squared coherence: this yielded a 415 × 415 functional association matrix each day, whose elements indicated the strength of functional connectivity between all pairs of nodes (FDR-thresholded at *q* < 0.05). Coherence offers several advantages over alternative methods for assessing connectivity: 1) estimation of *frequency-specific* covariances, 2) *simple interpretability* (values are normalized to the [0,1] interval), and 3) *robustness to temporal variability in hemodynamics* between brain regions, which can otherwise introduce time-lag confounds to connectivity estimates via Pearson correlation.

### Statistical analyses

Statistical analyses were conducted in MATLAB (version R2020b) and R (version 4.1.3).

#### Calculating time-synchronous variation in functional connectivity

First, we assessed time-synchronous variation in functional connectivity associated with testosterone, cortisol, and estradiol through a standardized regression analysis. Data were *Z*-transformed and edgewise coherence was regressed against hormonal timeseries to capture day-by-day variation in connectivity relative to hormonal fluctuations. For each model, we computed robust empirical null distributions of test statistics via 10,000 iterations of nonparametric permutation testing: under the null hypothesis of no temporal association between connectivity and hormones, the coherence data at each edge were randomly permuted, models were fit, and two-tailed *p*-values were obtained as the proportion of models in which the absolute value of the permuted test statistics equaled or exceeded the absolute value of the ‘true’ test statistics. We report edges surviving a threshold of *p* < .001. We did not apply additional corrections in an effort to maximize power in our small sample size.

As an additional sensitivity test to account for variability in wakefulness between morning and evening sessions, we included time since waking at each session (i.e. ∼1 hour since waking for morning sessions and ∼14 hours since waking for evening sessions) as a regressor in a supplemental analysis. Across all three steroid hormones, general trends of whole-brain coherence remain the same across networks, though mean nodal association strengths with estradiol and cortisol are slightly diminished (**Fig. S6**).

#### Determining network sensitivity to hormone fluctuations

For each time-synchronous model, we examined the direction of hormone-related associations and whether particular networks were more or less sensitive to hormonal fluctuations. Toward that end, we took the thresholded statistical parametric maps for each model (where edges are test statistics quantifying the magnitude of association between coherence and hormonal timeseries) and estimated nodal association strengths per graph theory’s treatment of signed, weighted networks. That is, positive and negative association strengths were computed independently for each of the 415 nodes by summing the suprathreshold positive/negative edges linked to them. We then assessed mean association strengths (±95% confidence intervals) in each direction across the various networks in our parcellation.

Finally, two-way ANOVAs with Tukey’s HSD (*p* < .05, corrected for family-wise error) was used to compare the variance in nodal association strengths among the three hormones (testosterone, cortisol, and estradiol) and all nine functional networks. Additional ANOVAs were used to compare the variance in nodal association strengths associated with testosterone and estradiol fluctuations among both datasets (male and female) and all networks.

## Results

### Endocrine assessments

As expected, hormone concentrations peaked in the morning and dipped in the evening (**Fig. 3**; **Table 1**). Testosterone, cortisol and estradiol were correlated to each other, and hormone concentrations from saliva samples (testosterone and cortisol) were tightly correlated with their serum sample counterparts (**Fig. S2; Table S1**). Testosterone is generally available in two forms: either bound to a carrier protein (sex hormone binding globulin (SHBG)) and therefore inert, or bioavailable (free or loosely bound to albumin). For completeness, this dataset provides assessments of both free and total testosterone. To maximize available MRI sessions, analyses involving testosterone and cortisol reflect values derived from salivary samples obtained across all 40 sessions. (**Fig. 1**). Morning to evening decreases in testosterone, estradiol, and cortisol were ∼63%, ∼39%, and ∼92%, respectively.

**Figure 3.**
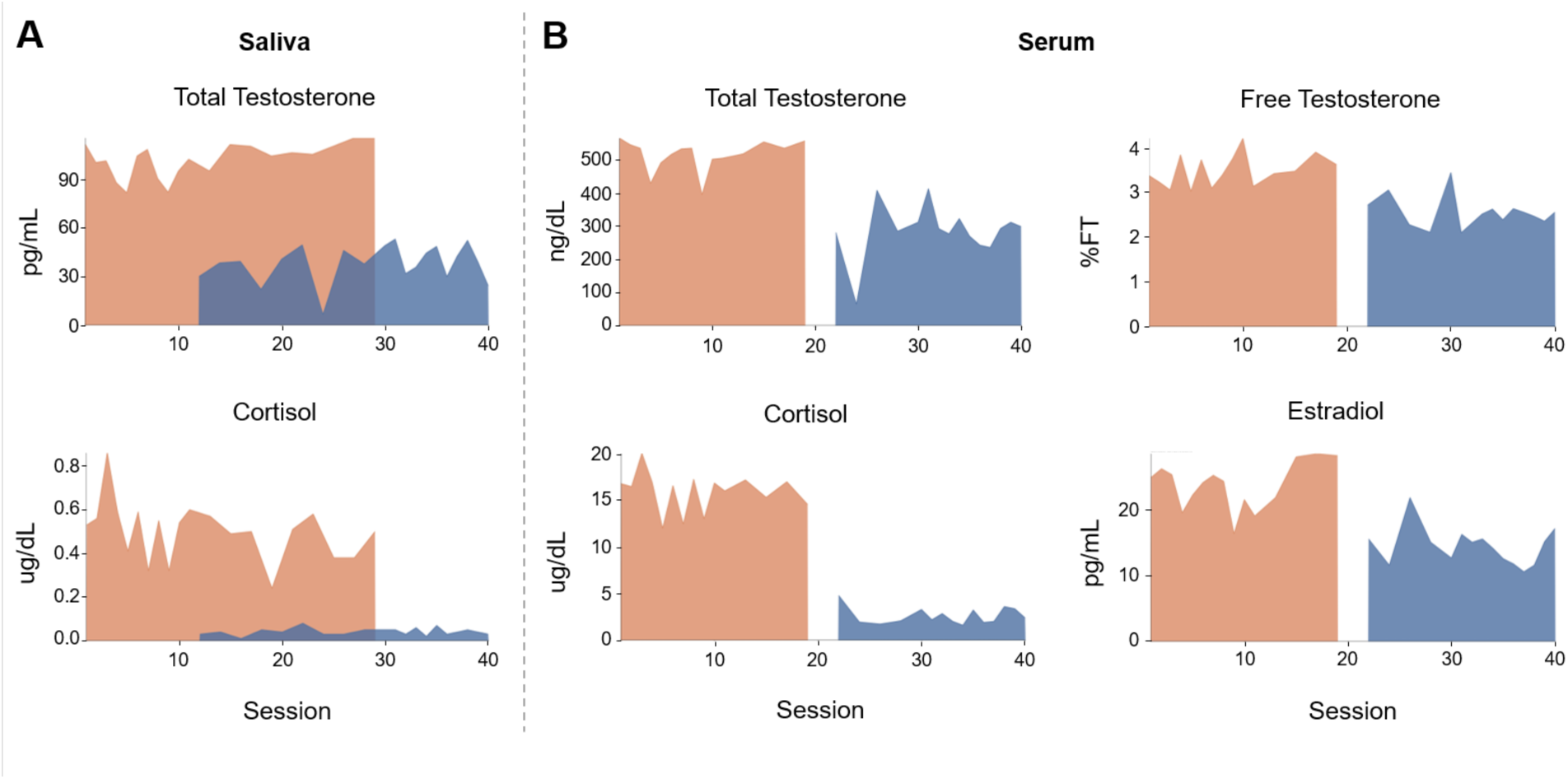
Steroid hormone concentrations by test session and time of day. **(A)** Saliva measurements for total testosterone and cortisol and **(B)** serum measurements for total testosterone, free testosterone, cortisol, and estradiol. All hormone concentrations were within or comparable to the standard range (see **Table 1**). Days 11-20 contained two sessions per day (sessions 11-30), with only one blood draw per day. On day 16 we switched the blood draws from the morning to the evening sessions, resulting in a two-session gap in serum values (i.e. sessions 20 and 21 do not have serum values).

**Table 1.** Gonadal hormones by time of day.

|  | Morning | Evening | Units |
| --- | --- | --- | --- |
|  | Mean (SD) | Mean (SD) |  |
| Total Testosterone (saliva) | 101.6 (10.2) | 37.9 (11.5) | pg/mL |
| Total Testosterone (serum) | 513.4 (47.0) | 286.4 (79.0) | ng/dL |
| Free Testosterone | 3.5 (0.4) | 2.6 (0.3) | %FT |
| Estradiol | 23.8 (3.6) | 14.5 (2.9) | pg/mL |
| Cortisol (saliva) | 0.5 (.1) | 0.04 (0.02) | ug/dL |
| Cortisol (serum) | 15.9 (2.1) | 2.6 (0.9) | ug/dL |

*Note.* Standard reference ranges based on aggregate data from Labcorp (https://www.labcorp.com/) for the serum values and ZRT Lab (https://www.zrtlab.com/) for saliva values. Standard hormone ranges are as follows: 41-141 pg/mL for total testosterone saliva samples, 264-916 ng/dL for total testosterone serum samples, 1.5-3.2% free testosterone samples, 7.6-42.6 pg/mL for estradiol samples, 0.37-0.95 ug/dL for morning cortisol saliva samples, 0.04-0.10 ug/dL for evening cortisol saliva samples, 6.2-19.4 ug/dL for morning cortisol serum samples, and 2.3-11.9 ug/dL for evening cortisol serum samples.

### Time-synchronous associations between steroid hormones and whole-brain functional connectivity

Previous work from our group (Pritschet et al., 2020) demonstrated robust increases in whole-brain coherence with increasing estradiol concentrations over the menstrual cycle in a naturally-cycling female. Here, we tested the hypothesis that whole-brain resting-state functional connectivity in a male is associated with diurnal intrinsic fluctuations in total testosterone, cortisol, and estradiol in a time-synchronous (i.e., session-to-session) manner. Based on previous findings, we predicted that the Default Mode, Dorsal Attention, and Temporal Parietal Networks would show the strongest associations with fluctuations in steroid hormone concentrations.

Increases in all three steroid hormones were associated with greater whole-brain functional connectivity across most of the 9 functional networks analyzed (**Fig. 4**). Notably, most networks showed some level of positive association strength on average (with most 95% CIs not intersecting zero). Efficiency (within-network integration) and participation (between-network integration) were not significantly different from morning to evening (*p* = .311 and *p* = .339, respectively) (**Fig. S3; Fig. S4**).

**Figure 4.**
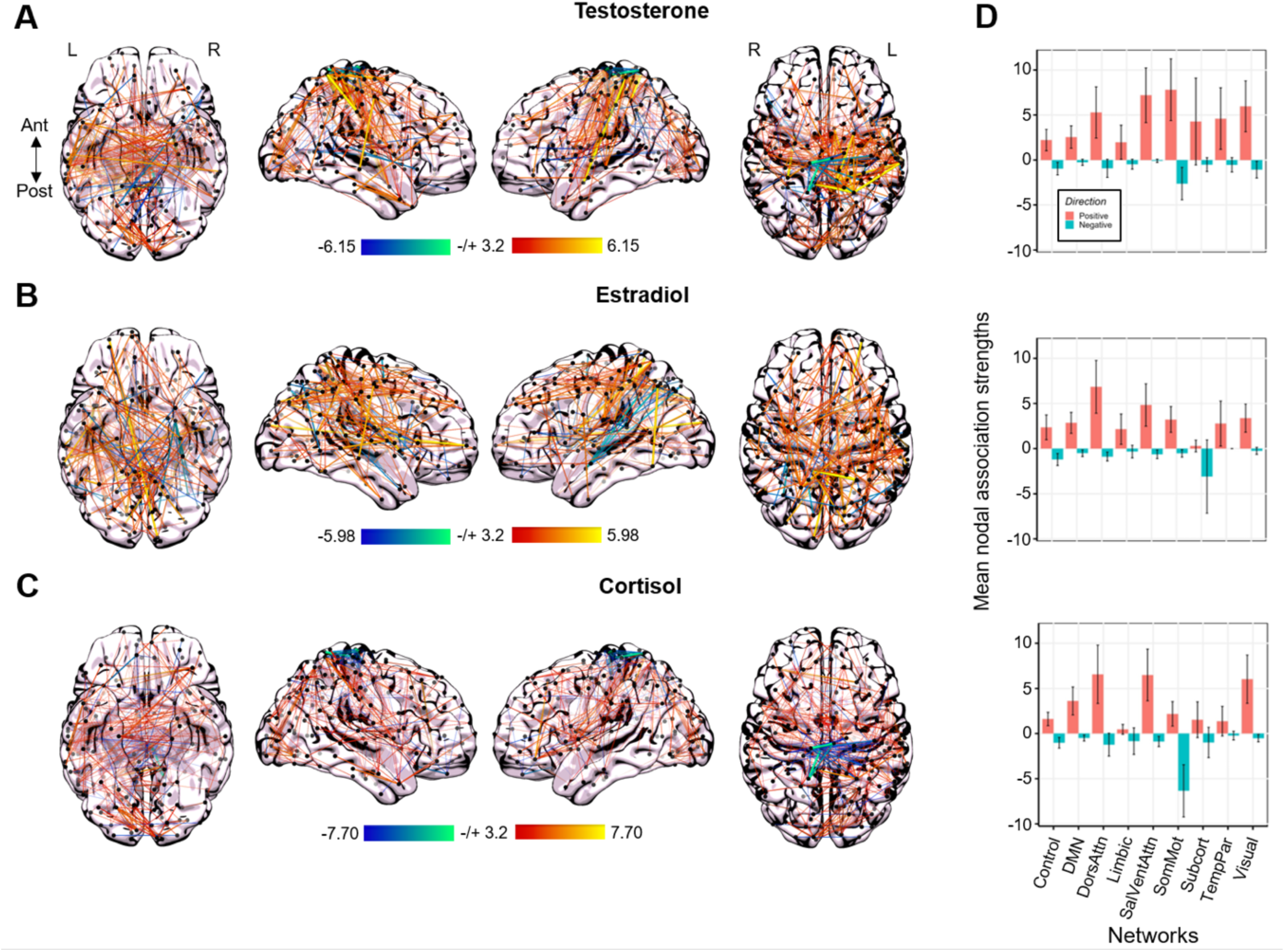
Whole-brain connectivity at rest is associated with intrinsic fluctuations in testosterone, cortisol, and estradiol. **(A)** Time-synchronous associations between total testosterone and coherence (left). Hotter colors indicate increased coherence with higher concentrations of testosterone; cool colors indicate the reverse. Reported edges survive a threshold of p<0.001. Mean nodal association strengths (right). Error bars give 95% confidence intervals. ‘Positive’ refers to the average magnitude of positive associations (e.g., stronger coherence with higher testosterone concentrations); ‘Negative’ refers to the average magnitude of inverse associations (e.g., weaker coherence with higher testosterone concentrations). Abbreviations: DMN, Default Mode Network; DorsAttn, Dorsal Attention Network; SalVentAttn, Salience/Ventral Attention Network; SomMot, SomatoMotor Network; TempPar, Temporal Parietal Network. **(B)** Time-synchronous associations between estradiol and coherence (left) and mean nodal association strengths (right). **(C)** Time-synchronous associations between cortisol and coherence (left) and mean nodal association strengths (right).

A two-way ANOVA revealed significant main effects of hormone (*F*(2,1218) = 3.352, *p* = .035) and functional network (*F*(8,1218) = 7.042, *p* < .001), and a significant interaction between hormone and network (*F*(16,1218) = 1.722, *p* = .037) for *positive* mean nodal association strengths. Tukey’s HSD post-hoc analyses revealed that testosterone-coherence associations were greater in magnitude than those observed for estradiol (*p* = .041). Additionally, the Dorsal Attention and Salience/Ventral Attention Networks showed the strongest positive associations with fluctuations across all three steroid hormones (**Fig. 4D**). Hormone-brain association strengths were greater in Dorsal Attention, Salience/Ventral Attention, and Visual Networks compared to the Control (*p* < .001, *p* < .001 and *p* = .011, respectively), and Limbic (*p* < .001, *p* < .001, and *p* = .033) Networks. Similarly, hormone-brain association strengths were greater in the Dorsal Attention and Salience/Ventral Attention Networks than the Default Mode (*p* = .002 and *p* = .002), and Subcortical (*p* = .034 and *p* = .039) Networks.

A separate analysis examining *negative* mean nodal association strengths showed statistically significant main effects of hormone (*F*(2,1218) = 7.524, *p* < .001) and functional network (*F*(8,1218) = 9.182, *p* < .001), and a significant interaction of hormones and networks (*F*(16,1218) = 4.338, *p* < .001). Tukey’s post-hoc analyses revealed that cortisol was associated with a greater magnitude of negative whole-brain coherence associations compared to testosterone (*p* = .012) and estradiol (*p* < .001). Across all hormones, the SomatoMotor Network was unique in showing strong negative mean nodal association strengths compared to all other networks, including Control (*p* < .001), Default Mode (*p* < .001), Dorsal Attention (*p* < .001), Salience/Ventral Attention (*p* < .001), Temporal Parietal (*p* < .001), Limbic (*p* < .001), and Visual (*p* < .001) Networks (**Fig. 4D**).

### Whole-brain coherence tied to intrinsic fluctuations of sex hormones in both sexes

To understand differences in the extent to which male and female brains respond to varying fluctuations in sex hormones, we compared whole-brain patterns of estradiol- and testosterone-related effects in our male participant to data from a previous dense-sampling study in a naturally-cycling female (**Fig. 5**) collected under a near-identical protocol (Pritschet et al., 2020). Notably, the overall positive associations between sex hormones and whole-brain coherence were robust across both sexes (**Fig. 5A and 5B**). In both participants, nearly all networks displayed strong positive associations with estradiol (the sole exception was the Subcortical Network in the male participant). Similarly, network strengths were positively associated with testosterone for both participants, and the magnitude of this relationship was heightened in the male participant. As a reference, the male participant experienced morning-to-evening decreases in testosterone, estradiol, and cortisol (∼63%, ∼39%, and ∼92%, respectively). The naturally cycling female participant experienced a 92% decrease in estradiol concentrations from ovulation to menses, and a 40% decrease in testosterone concentrations from the peak during the luteal phase to menses.

**Figure 5.**
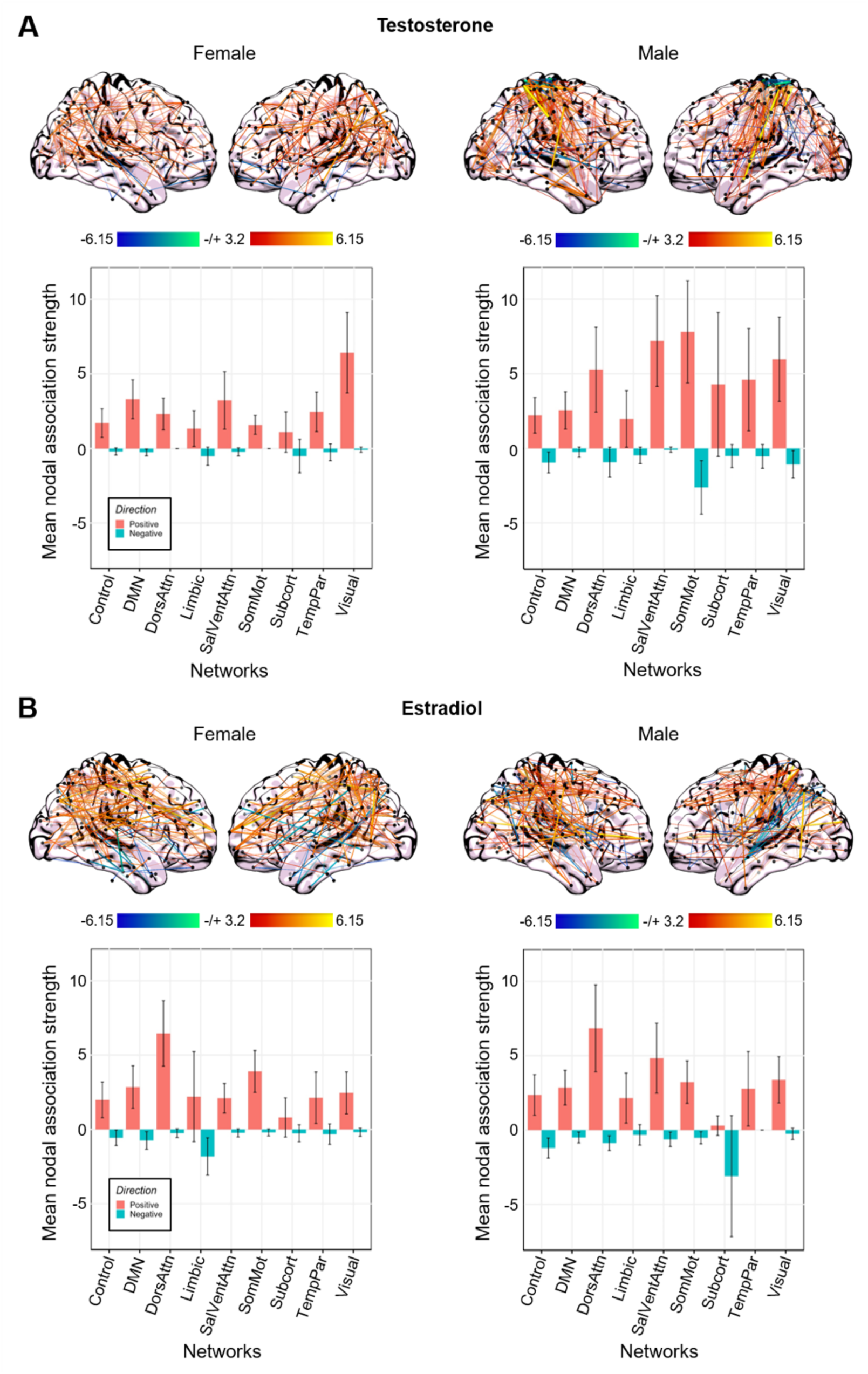
Intrinsic fluctuations in sex steroid hormones are associated with whole-brain resting-state functional connectivity to a comparable degree in a male and a naturally-cycling female. **(A)** Time-synchronous associations (top) and mean nodal association strengths (bottom) between total testosterone and coherence in a female (left) and male (right). **(B)** Time-synchronous associations (top) and mean nodal association strengths (bottom) between estradiol and coherence in a female (left) and male (right).

A two-way ANOVA revealed a statistically significant main effect of network (*F*(8,812) = 3.781, *p* < .001) and sex (*F*(1,812) = 15.635, *p* < .001), and a significant interaction between sex and network (*F*(8,812) = 2.961, *p* = .003), on positive mean nodal association strengths of whole-brain coherence associated with total testosterone fluctuations. Positive mean nodal association strengths were significantly greater in the male participant. Tukey’s post-hoc analyses revealed that fluctuations in whole-brain coherence correlated with testosterone concentrations in the Visual Network were significantly greater in magnitude than the Control Network (*p*=0.001), Default Mode (*p* = .020), and Limbic (*p* = .016) Networks. Positive strengths associated with testosterone fluctuations in the Salience/Ventral Attention Network were also significantly greater than strengths in the Control Network (*p* = .029). In the female participant, the strongest associations with testosterone were in the Visual, Default Mode, and Salience/Ventral Attention Networks. In the male participant, the strongest associations with testosterone were in the SomatoMotor, Salience/Ventral Attention, Dorsal Attention, and Visual Networks (**Fig. 5A**). When comparing equivalent networks in both sexes, the SomatoMotor Network was the only network to reach a significant difference (*p* < .001).

A two-way ANOVA revealed a statistically significant main effect of network (*F*(8,812) = 5.940, *p* < .001), but not sex (*F*(1,812) = 1.081, *p* = .299), on positive mean nodal association strengths of whole-brain coherence associated with estradiol fluctuations. Additionally, there was no significant interaction between sex and network (*F*(8,812) = .709, *p* = .684), with networks displaying comparable magnitudes of association strengths in both sexes. Tukey’s post-hoc analyses revealed that fluctuations in whole-brain coherence positively correlated with estradiol concentrations in the Dorsal Attention Network were significantly greater in magnitude than every other network: Control (*p* < .001), Default Mode (*p* < .001), Salience/Ventral Attention (*p* = .004), Temporal Parietal (*p* = .015), Limbic (*p* < .001), SomatoMotor (*p* = .002), Subcortical (*p* < .001), and Visual (*p* < .001) Networks (**Fig. 5B**).

#### Nodes most strongly tied to fluctuation in steroid hormone vary across the sexes

Within the male participant, nodes in the Dorsal Attention, Salience/Ventral Attention, and Visual Networks demonstrated the strongest association strengths with diurnal variation in all three steroid hormones. Nodes in the bilateral postcentral region and right superior parietal lobe (Dorsal Attention Network), right parietal operculum, and right insula (Salience/Ventral Attention Network), and bilateral regions of the SomatoMotor Network showed the greatest associations between functional activation and diurnal variation in testosterone, estradiol, and cortisol concentrations. Further, bilateral regions of the extrastriate cortex (Visual Network) and right anterior temporal lobe (Default Mode Network) were strongly tied to fluctuations in testosterone and cortisol concentrations, but *not* to fluctuations in estradiol concentrations. Regions in the right lateral PFC and right precuneus (Control Network) were most strongly tied to diurnal fluctuations of estradiol and testosterone concentrations.

The nodes tied most strongly to estradiol fluctuations in the female participant included the left superior parietal lobe and postcentral region (Dorsal Attention Network) and bilateral regions of the somatomotor cortex (SomatoMotor Network), while the left posterior cingulate cortex and left dorsal PFC (Default Mode Network) and bilateral superior extrastriate cortex (Visual Network) were most strongly tied to fluctuations in testosterone.

## Discussion

This precision imaging study reveals rhythmic changes in the brain’s functional network architecture tied to diurnal fluctuations in steroid hormones. In a previous study, we mapped the brain’s response to intrinsic fluctuations in sex hormones across the menstrual cycle in a densely-sampled female (Pritschet et al., 2021, 2020; Fitzgerald et al., 2020; Mueller et al., 2021; Taylor et al., 2020). Here, we extend those findings by densely sampling a male with MRI and venipuncture every 12-24h, capturing the brain’s response to diurnal changes in steroid hormone secretion from AM to PM. Fluctuations in testosterone, cortisol, and estradiol were tied to changes in functional coherence across most of cortex. In particular, Dorsal Attention and Salience/Ventral Attention Networks showed strong associations across all three hormones.

Comparisons between the densely-sampled male and female revealed estradiol’s ability to influence whole-brain coherence, regardless of sex. Similarly, testosterone fluctuations influence intrinsic network properties in both sexes with exaggerated effects in the male, demonstrating that sex hormones impact brain function in both sexes, and these effects are not limited to the menstrual cycle.

Existing evidence supports an association between time-of-day and functional connectivity. Sparse-sampling studies investigating the impact of diurnal variation on brain dynamics tested subjects at 2+ timepoints (e.g. one morning and one evening session) and compared brain metrics across these timepoints, demonstrating a time-of-day effect on functional brain metrics (e.g., small-worldness, assortativity, and global signal amplitude; Farahani et al., 2022; Orban et al., 2020; Shannon et al., 2013; Marek et al., 2010). Consistent with our findings, Farahani et al. (2022) found that the ventral attention and visual networks are enhanced during morning sessions compared to evening sessions. Our dense-sampling study, which tracks dynamic changes in the brain over time with super-high temporal precision, sheds further light on these findings by probing a specific biological driver of time-of-day effects in network connectivity: namely, diurnal steroid hormone fluctuations.

Though we cannot test *how* steroid hormones are indirectly influencing network dynamics through cellular action, we know from rodent models that steroid hormones induce rapid effects on cell morphology. In rats, hormone-induced changes in synaptic plasticity have been studied extensively in hippocampal CA1 neurons, where fluctuations in androgen and estrogen concentrations are correlated with dendritic spine density (Hojo et al., 2008; Leranth et al., 2003, 2004; Li et al. 2012; MacLusky et al., 2006; Tozzi et al., 2019). Our data suggests that rapid effects of steroid hormones are also observable at the macroscopic level of intrinsic functional networks in the human brain. The A.M. diurnal peak in steroid hormones is tied to increases in whole-brain coherence, particularly Dorsal Attention, Salience/Ventral Attention, and Visual Networks.

Direct comparisons between a male and female participant debunk a persistent myth that sex steroid hormones matter more for the ‘female brain’ than the ‘male brain’. Here we show that intrinsic fluctuations in steroid hormones – be it across the menstrual cycle or diurnal cycle – modulate cortical activity in the human brain and the magnitude of these effects is comparable across the sexes. Dynamic changes in estradiol production impacted brain networks to a similar degree in a densely sampled male and female. Notably, we see this effect despite significantly different estradiol concentrations (*t*(29.64) = -6.39, *p* < .001) in the male participant and the female participant. Further, diurnal changes in testosterone had a greater impact on brain networks in the male participant than those observed in the female participant across the cycle.

Though similar networks (i.e. the Dorsal Attention, SomatoMotor, Default Mode, and Visual Networks) demonstrated associations with fluctuations in sex hormones across the male and female participant, different nodes within these networks were most strongly tied to sex hormone fluctuations with exception of the left postcentral region of the Dorsal Attention Network, which was tied to fluctuations in estradiol concentrations in both sexes. Both testosterone and estradiol are neuroprotective and age-related decreases in concentrations of both sex hormones are associated with cognitive decline (Corona et al., 2021; Hogervorst et al., 2004; Holland et al., 2011; Janowsky, 2006; Luine, 2014; Wolf & Kirschbaum, 2001), therefore it is not surprising that estradiol may have an analogous effect on neural activity at rest in both sexes. Importantly, this data challenges the notion that estradiol and testosterone are female- and male-specific hormones, respectively—both hormones influence brain function regardless of sex.

Diurnal rhythms are ubiquitous across mammalian species and an organizing principle for human physiology and behavior. This precision imaging study sheds light on the extent to which intrinsic rhythms in hormone production influence large scale brain networks by collecting fluid biomarkers and MRI data on a participant at an unprecedented temporal frequency. However, one limiting factor of this single subject study is the extent to which the observed findings generalize to the broader population. Behavioral factors that vary by time of day, including sleep quality, stress, and aggression, may contribute to the observed changes in brain network dynamics. It is unlikely that these factors could fully explain our observed results given that the participant maintained a consistent and low level of stress and aggression and maintained a stable sleep schedule throughout the duration of the study. Future dense-sampling studies ought to investigate the extent to which these observations are universal vs. idiosyncratic across a diverse range of individuals. The magnitude of diurnal variation in hormone concentrations diminishes with advanced age, so variation in whole-brain coherence from morning to evening may weaken with age as well, a topic future studies could investigate.

Additionally, other physiological factors such as alertness, respiratory rate, global changes in blood flow, or other endocrine changes such as melatonin may contribute to our observed results. Orban et al. (2020) found that the magnitude of fluctuations in global brain activity decline throughout the day. We factored in global activity into our analysis, producing an overall decrease in mean nodal association strengths though the overall pattern of strengths across all networks remained the same, suggesting that global activity can account for some of the diurnal variation within networks. Moreover, melatonin production and individual sleep-wake preference (i.e. individuals with earlier vs. later sleep and wake times) has been shown to affect functional connectivity within the Default Mode Network, with “night owl” subjects displaying reduced functional connectivity during typical work hours (Facer-Childs et al., 2019). While our precision imaging study avoided these confounds (the subject displayed a highly metronomic sleep/wake cycle over the 40 day experiment), future studies examining diurnal rhythms in an expanded sample should control for individual sleep-wake preferences. In a supplemental analysis, we included total awake time at each session as a regressor to control for potential differences due to alertness (**Fig. S6**), and the pattern of results did not change appreciably.

This dataset provides a novel approach for understanding endocrine modulation of functional networks and provides a starting point for future studies to probe the functional significance of these rhythms for behavior. Combined with our findings in a naturally-cycling female, these results further demonstrate steroid hormones’ ability to modulate brain function across both sexes, and further research in this field is critical for expanding our basic understanding of endocrine influences on the brain.

### Author Contributions

Conceptualization: L.P, P.S., E.G.J.; Formal analysis: H.G.; Funding Acquisition: E.G.J.; Investigation: L.P., P.S; Methodology: L.P., T.S.; Project Administration: H.G., E.M., L.P.; Resources: E.G.J.; Supervision: E.G.J.; Visualization: H.G., E.M; Writing – Original Draft: H.G.; Writing – Review and Editing: L.P, P.S., T.S., E.M., E.G.J.

## Acknowledgements

This study was supported by the Ann S. Bowers Women’s Brain Health Initiative (EGJ), UC Academic Senate (EGJ), and NIH AG063843 (EGJ). We thank Mario Mendoza for phlebotomy and MRI assistance.

