## Supplemental Figures for "Diurnal fluctuations in steroid hormones tied to variation in intrinsic functional connectivity in a densely sampled male"

### Supplemental Tables

Supplementary Table 1. Correlations between gonadal hormones.

| Hormone Pair | | t-statistic | p-value | Pearson's correlation |
|---|---|---|---|---|
| Cortisol (serum) | Cortisol (saliva) | 19.98 | 2.2e-16*** | 0.97 |
| Total Testosterone (serum) | Total Testosterone (saliva) | 15.22 | 4.6e-15*** | 0.94 |
| Total Testosterone (serum) | Free Testosterone | 3.92 | 5.2e-04** | 0.60 |
| Total Testosterone (saliva) | Free Testosterone | 5.65 | 4.7e-06*** | 0.73 |
| Cortisol (serum) | Total Testosterone (serum) | 9.24 | 5.3e-10*** | 0.87 |
| Cortisol (saliva) | Total Testosterone (saliva) | 11.38 | 8.4e-14*** | 0.88 |
| Total Testosterone (serum) | Estradiol | 11.83 | 2.1e-12*** | 0.91 |
| Total Testosterone (saliva) | Estradiol | 8.88 | 1.2e-09*** | 0.86 |
| Cortisol (serum) | Estradiol | 7.57 | 3.0e-08*** | 0.82 |
| Cortisol (saliva) | Estradiol | 6.05 | 1.6e-06*** | 0.75 |

*Note.* Bonferroni adjusted alpha for 15 comparisons: *p<0.003, **p<0.0007, ***p<0.00007

#### Supplemental Figures

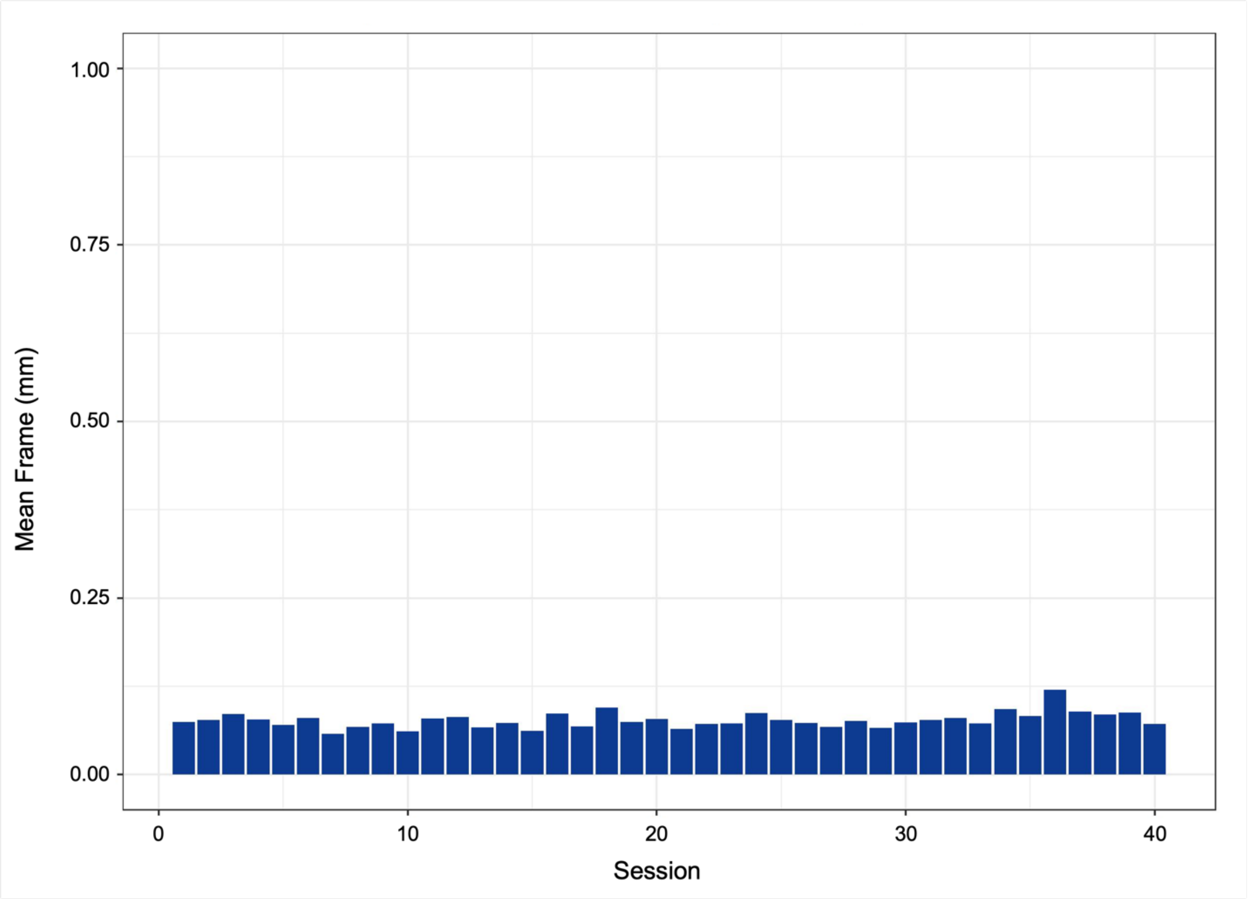


**Supplementary Figure 1. Mean framewise displacement measurement (mm) averaged across each session's resting state scan.** Head motion was minimal compared to a conservative 'acceptable' limit of 1mm.

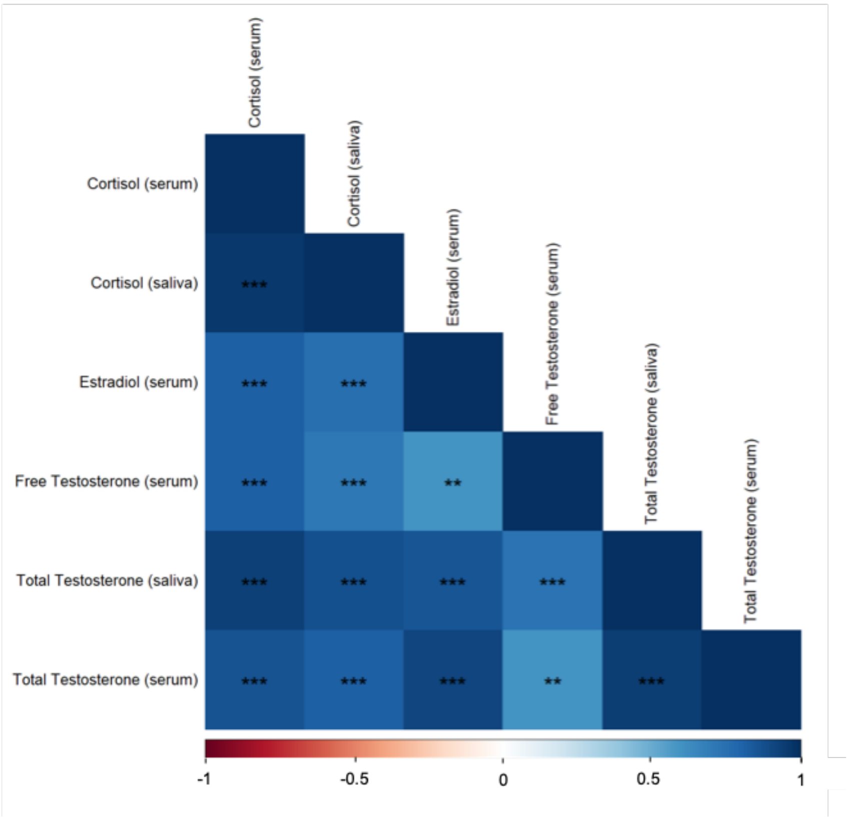


**Supplementary Figure 2. All steroid hormone concentrations were tightly correlated with each other.** All saliva samples were tightly correlated with their serum sample counterparts. Asterisks indicate significant correlations after Bonferroni correction (* $p < 0.003$, ** $p < 0.0007$, *** $p < 0.00007$).

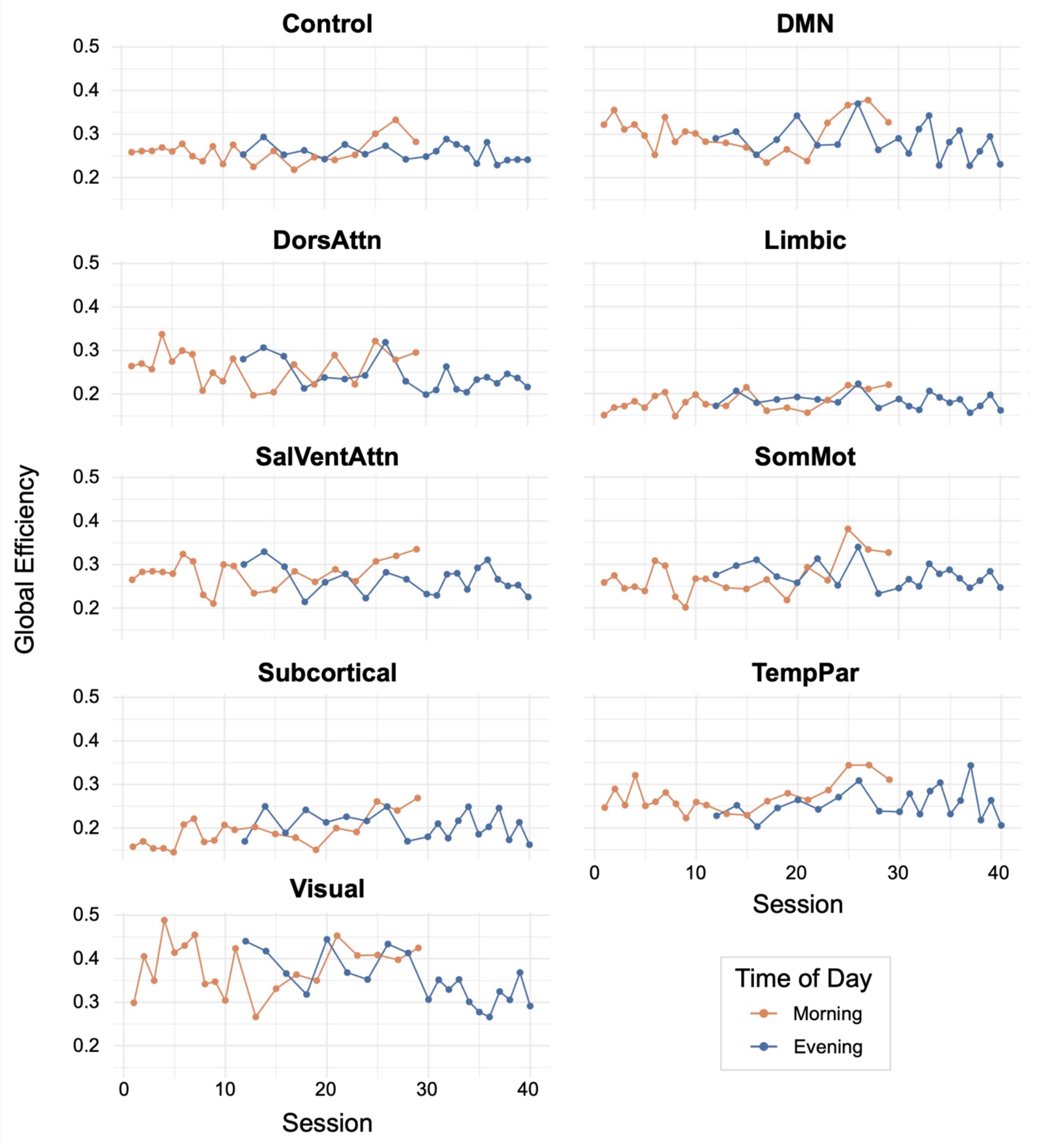


**Supplemental Figure 3. Efficiency values at each session by time of day.** Efficiency was not significantly different from morning to evening.

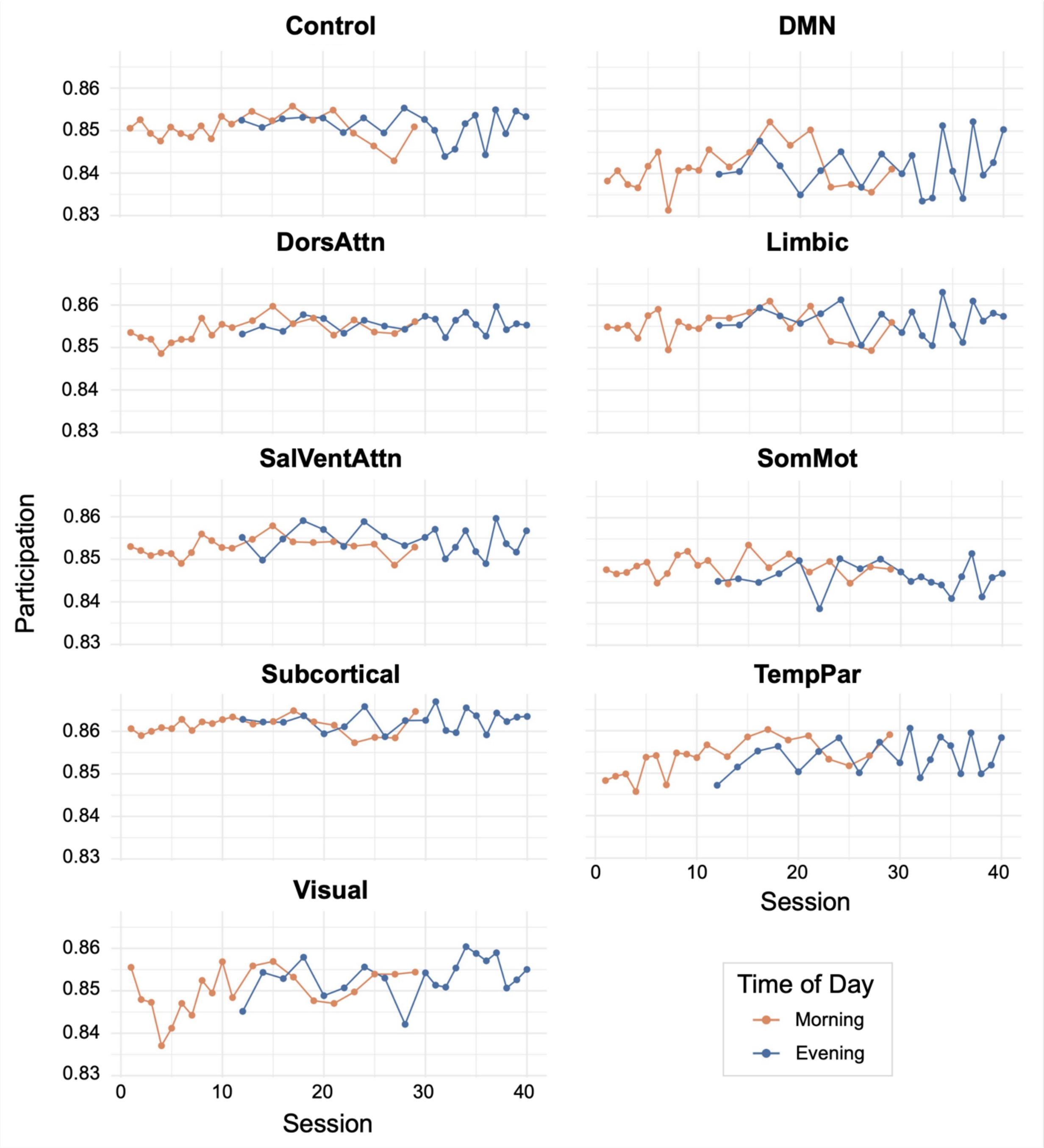


**Supplemental Figure 4. Participation values at each session by time of day.** Participation was not significantly different from morning to evening.

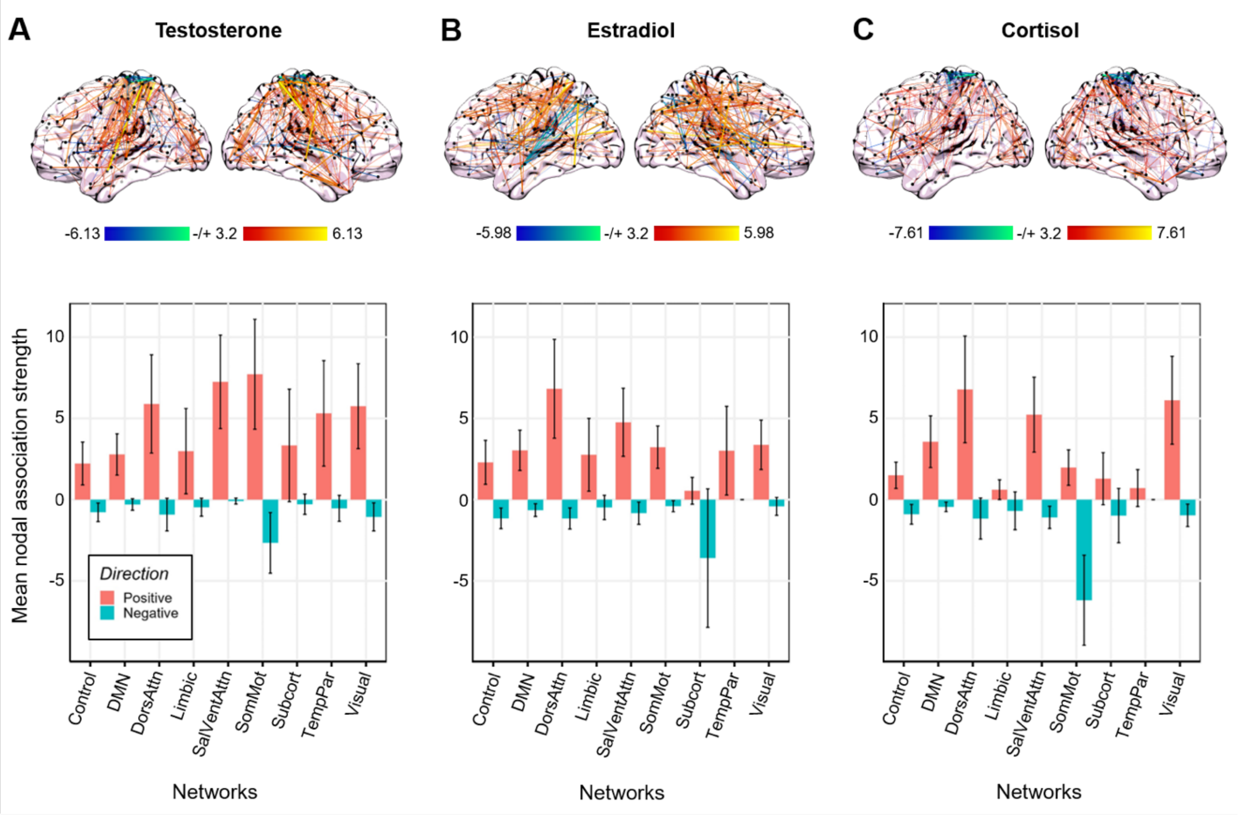


**Supplementary Figure 5. Time-synchronous brain-hormone associations without global signal regression.** While the average magnitude of brain-hormone associations differed in some networks, the overall trends remained. **(A)** Time-synchronous associations between total testosterone and coherence (top) and mean nodal association strengths by network (bottom). Hotter colors indicate increased coherence with higher concentrations of testosterone; cool colors indicate the reverse. Reported edges survive a threshold of $p<0.001$. **(B)** Time-synchronous associations between estradiol and coherence (top) and mean nodal association strengths by network (bottom). **(C)** Time-synchronous associations between cortisol and coherence (top) and mean nodal association strengths by network (bottom).

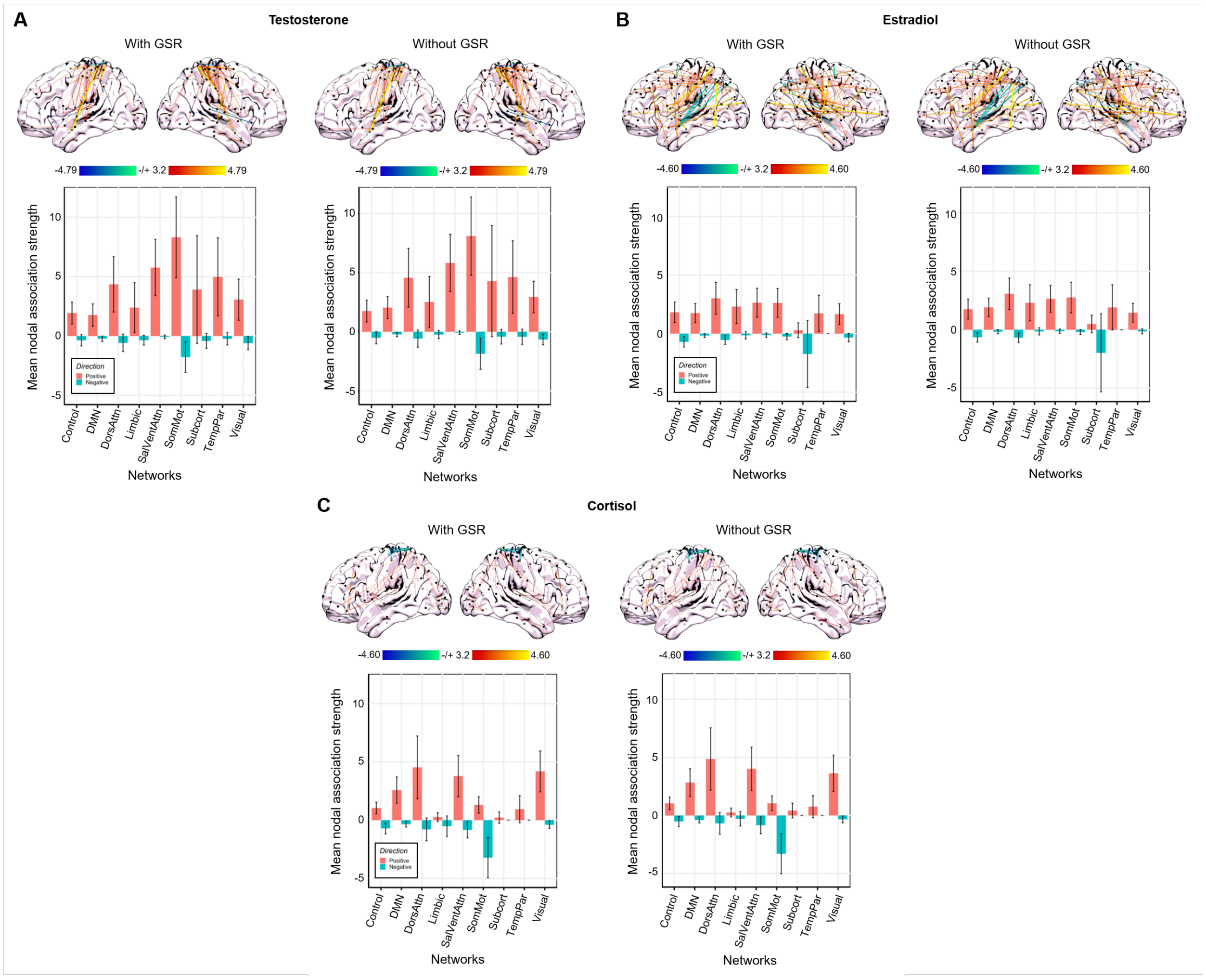


**Supplementary Figure 6. Time-synchronous brain-hormone associations accounting for awake time.** When incorporating time awake at each scan as a regressor, the average magnitude of brain-hormone associations differed in some networks, though overall trends remained and were comparable with and without global signal regression. Time-synchronous associations between total testosterone **(A)**, estradiol **(B)**, and cortisol **(C)** and coherence, and mean nodal association strengths by network (bottom). Data is presented with (left) and without (right) global signal regression.
